# PTEN inhibits AMPK to control collective migration

**DOI:** 10.1101/2021.12.13.472380

**Authors:** Florent Peglion, Lavinia Capuana, Isabelle Perfettini, Ben Braithwaite, Emie Quissac, Karin Forsberg-Nilsson, Flora Llense, Sandrine Etienne-Manneville

## Abstract

PTEN is one of the most frequently mutated tumor suppressor gene in cancer. PTEN is generally altered in invasive cancers such as glioblastomas, but its function in collective cell migration and invasion is not fully characterized. Herein, we report that the loss of PTEN increases cell speed during collective migration of non-tumourous cells both in vitro and in vivo. We further show that loss of PTEN promotes LKB1-dependent phosphorylation and activation of the major metabolic regulator AMPK. In turn AMPK increases VASP phosphorylation, reduces VASP localization at cell-cell junctions and decreases the interjunctional transverse actin arcs at the leading front, provoking a weakening of cell-cell contacts and increasing migration speed. Targeting AMPK activity not only slows down PTEN-depleted cells, it also limits PTEN-null glioblastoma cell invasion, opening new opportunities to treat glioblastoma lethal invasiveness.

## INTRODUCTION

At the heart of a myriad of cellular processes, PTEN (Phosphatase and TENsin homolog deleted in chromosome ten) is one of the most altered tumour suppressors in human cancer ^1, 2^. This holds particularly true in glioblastoma (GBM), the most malignant and frequent brain tumour, where PTEN alteration is observed in 41% of cases ^3-5^. PTEN is a dual-specific protein and lipid phosphatase ^6, 7^. By dephosphorylating phosphatidylinositol-3, 4, 5-triphosphate (PIP_3_) into phosphatidylinositol-4, 5-bisphosphate (PIP_2_), PTEN antagonizes the pro-oncogenic PI3K-Akt signalling pathway ^8, 9^ which is key to coordinate cell proliferation, growth, survival and metabolism. PTEN protein-phosphatase activity also plays a role in PTEN functions but molecular details remain scarce ^10^. Cancer cells are characterized by increased invasive capacities, which facilitate cancer spreading, and metastases. PTEN ability to regulate PIP_3_/PIP_2_ level at the plasma membrane enables it to control cell polarization and directionality during the directed migration of single cells ^11-13^. However, cancer invasion often involves collective motility ^14^ and the role of PTEN in the control of collective migration and invasion is still unclear. Since PIP_3_ is a key determinant of the basolateral surface, reduced PTEN activity has been proposed to alter epithelial characteristics, causing cells to switch to an invasive, motile, mesenchymal phenotype ^15^. PTEN rescue experiments in cancer cell lines highlighted the importance of lipid phosphatase-independent activities, in particular in GBM cells ^16-18^. In NIH 3T3 cells and U-87 GBM cells, PTEN overexpression was shown to decrease cell migration and invasion possibly by reducing tyrosine (Y) phosphorylation of focal adhesion kinase (FAK) ^16, 19^. However, it is still unclear how PTEN loss alone can promote cell migration and invasion of tumour cells.

To tackle this question, we down-regulated endogenous PTEN both in primary glial cells *in vitro* and in endothelial cells *in vivo* and demonstrated that, during collective migration, PTEN depletion increases the speed of migrating cells. We unravel a new lipid-phosphatase independent connection between PTEN and the bioenergetics master regulator AMPK, that controls actin remodelling and cell-cell junctions to maintain cohesion and keep collective glial cell migration and invasion in check.

## RESULTS

To address the effects of PTEN loss in collective migration we designed siRNA against PTEN to decrease PTEN expression in rat astrocytes. siRNA efficiency was validated by observing a >90% decrease of total PTEN level and a sharp increase of PTEN-opposing PI3K pathway activity, highlighted by upregulation of pAKT level (Supplementary Figure 1A). Collective migration of glial cells was assessed using an in vitro wound-healing assay, which allows the quantitative assessment of cell speed and polarity^20^. Compared to control astrocytes (siCTL), PTEN-depleted cells (siPTEN) closed the artificial wound significantly faster (Figure 1A). The analysis of single cell tracks showed that PTEN-depleted wound-edge cells migrate longer distance than control cells (Figure 1B). Quantification revealed that PTEN loss strongly increases cell velocity (+70% Figure 1C) without strong defects in directionality and persistence of direction (Supplementary Figure 1B,C). Expression of a GFP-tagged wild-type PTEN construct, but not of a phosphatase dead mutant C124S (Supplementary Figure 1D), significantly slows down the migration of siPTEN astrocytes (Figure 1B and 1C). Taken together these data reveal that deletion of PTEN is sufficient to increase cell speed during collective cell migration.

**Figure 1:**
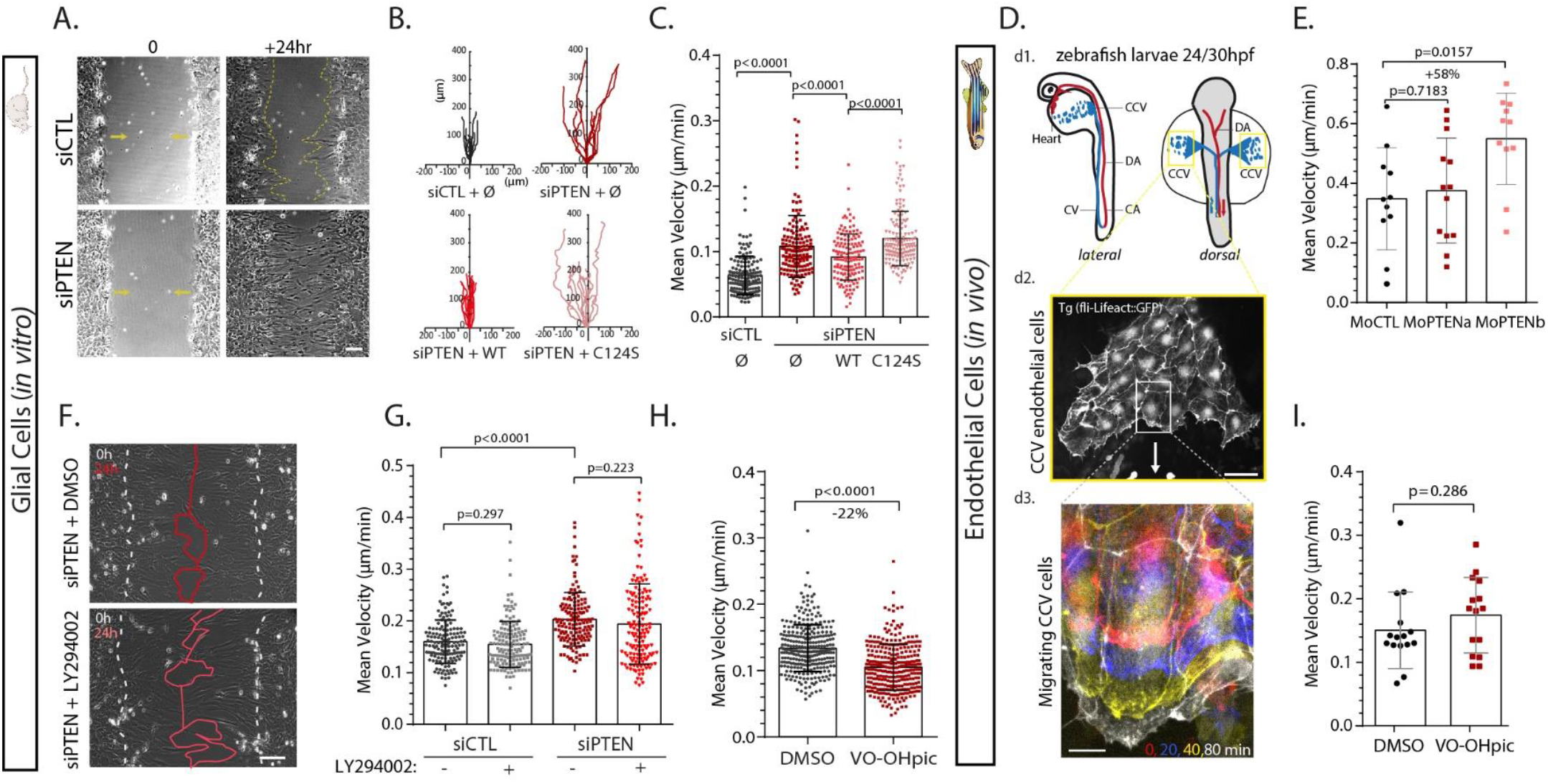
PTEN loss enhances collective cell migration independently of PI3K/AKT signalling. **A**. Phase-contrast images of control (siCTL) and PTEN-depleted (siPTEN) astrocytes migrating collectively in a wound-healing assay. **B**. Diagrams showing trajectories covered by 10 representative siCTL, and siPTEN cells rescued with GFP alone (Ø), GFP-PTENwt (WT) or phosphatase dead GFP-PTENC124S (C124S), in 24hours. **C**. Mean velocity of the same cells (n=142, N=3, ANOVA analysis). **D**. *d1*. Schemes representing lateral and dorsal view of a 24/30hpf zebrafish embryo with dorsal (DA) and caudal arteries (CA) in red and the caudal (CV) and common cardinal (CCV) veins. CCV endothelial cells are delaminating from the midline, moving towards the heart (yellow box). *d2*. Representative fluorescent image of migrating CCV cells expressing LifeAct::GFP. Scale bar: 50µm. *d3*.Time-colored zoomed-in (white rectangle, *d2*) image of migrating cells. Red is t=0, blue is t=20min, yellow is t=40min and white is t=180min. Scale bar represents 10µm. **E**. Mean velocity of CCV endothelial cells in control zebrafish embryos (MoCTL), PTENa morphant (MoPTENa) and PTENb morphant (MoPTENb) (n=11,13,12; N=3, Mann-Whittney analysis). **F**. Phase-contrast images of PTEN-depleted cells treated with DMSO or the PI3K inhibitor LY294002. White dashed lines delineate the border of the wound at t=0. Red lines delineate the border of the monolayer 24h later. Scale bar A. and D.:100µm. **G**. Mean velocity of siCTL and siPTEN cells treated with or without LY294002 (n=198 cells, N=3, ANOVA analysis). **H**. Mean velocity of astrocytes treated with DMSO or VO-OHpic (n=300, N=3, unpaired Student t-test analysis). **I**. Mean velocity of CCV endothelial cells in zebrafish embryos treated with DMSO or VO-OHpic (n=15, N=3, Mann-Whittney analysis).

To assess whether PTEN function in collective migration was conserved *in vivo*, we looked at the endothelial cells (EC) forming the communal cardinal vein (CCV) during zebrafish (*Danio Rerio*) early development. Starting around 20hr post fecundation, ECs delaminate from the midline and move collectively towards the heart, between the epidermis on top and the yolk sac syncytium layer underneath (Figure 1d1). ECs migrate as a single-sheet monolayer (Figure 1d2), in a cadherin-dependent fashion ^21^, similarly to astrocyte monolayers closing an in vitro artificial wound, despite being much faster (Figure 1d3). A consequence of whole-genome duplication in teleosts, the zebrafish genome encodes two *pten* genes, *ptena* and *ptenb*, ubiquitously expressed and sharing partially redundant functions during early development ^22, 23^. Using morpholinos specific to *pten* orthologs, we observed that in *ptenb* morphants, but not in *ptena* morphants, ECs migrate faster than in control morphant fish (+58% increase) (Figure 1E). *Ptena* may not be significantly expressed in ECs of the CCV at this stage, or it does not contribute to collective cell migration. In fact, *ptena* and *<u>ptenb</u>* both share an identical phosphatase domain to human PTEN but differ by their membrane localisation C2 motif ^22^. A different subcellular localisation could thus explain divergent roles of the two isoforms. Altogether, our results show that PTEN limits cell speed during collective migration in different cell types and models.

PTEN lipid phosphatase activity, which antognizes PI3K activity, is crucial to establish a front-rear gradient of lipids that sustain chemotaxis and directionality of amoeba and immune single cell migration^12^. To test the role of phospholipid signalling in the migration of PTEN-depleted cells, glial cells were treated with PI3K inhibitor LY294002. LY294002 treatment, whose efficiency is supported by a 80% drop in pAKT/AKT level compared to DMSO in control (Supplementary Figure 1E,F) and siPTEN cells (Supplementary Figure 1F), does not affect the speed of migration of PTEN-depleted cells (Figure 1F,G). Additionally, treatment with VO-OHpic, a potent inhibitor of PTEN lipid phosphatase activity that does not block protein phosphatase action of PTEN-like CX5R motif bearing phosphatase PTP1B ^24, 25^ strongly increases AKT phosphorylation (Supplementary Figure 1E) but does not increase astrocyte velocity (Figure 1H). Similarly, VO-OHpic addition to zebrafish embryos does not significantly alter mean ECs migration velocity (Figure 1I). Taken together, these data show that the increased cell velocity observed following PTEN depletion is independent of the PI3K/AKT pathway.

Collective cell migration relies on the synchronization of pathways permitting cytoskeleton remodelling and the maintenance of intercellular adhesion ^26-28^. To unveil how PTEN loss leads to an increase in cell migration velocity, we investigated the impact of PTEN depletion on the actin cytoskeleton and the cell-cell junctions. In the front row of migrating astrocytes, F-actin form both longitudinal fibres anchored at the leading edge focal adhesions and Interjunctionnal Transverse Arcs (ITA) that are oriented perpendicularly to the direction of migration and connect neighbouring cells through adherens junctions ^29^ (Figure 2A, D). Interestingly, PTEN depletion leads to a loss of ITAs (Figure 2A, C, D) while actin cables parallel to the direction of movement become more pronounced as shown by the changes in the distribution of actin cables orientation within the cell (Figure 2B). *In vivo*, front row of migrating ECs from control zebrafish morphants expressing lifeActGFP commonly show similar ITA (Figure 2E). Depletion of *ptenb* decreased the number of cells connected by ITA (Figure 2E, F). In parallel with the perturbation of ITAs, PTEN depletion also decreased cell-cell junction’s linearity in both glial cells *in vitro* and ECs *in vivo* (Figure 2G, H). Altogether, these data show PTEN regulates actin organization at cell-cell contacts to support connectivity between leader cells during collective migration (Figure 2I).

**Figure 2:**
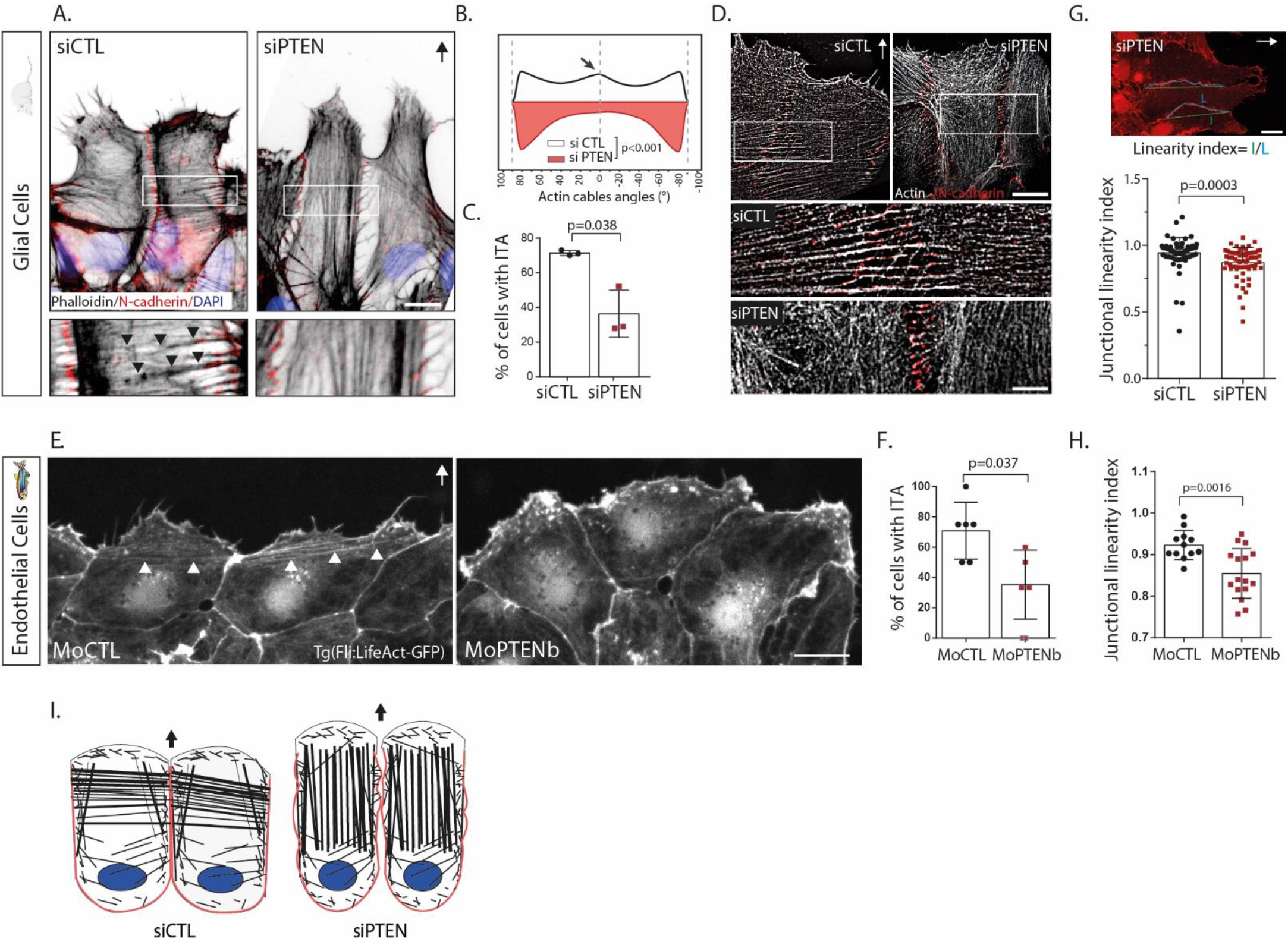
PTEN controls first-row cell-cell cohesion. **A**. Immunofluorescence images of actin filaments and cell-cell junctions in siCTL and siPTEN migrating astrocytes stained with phalloidin (black), N-cadherin (red) and Dapi (blue). White arrow, direction of migration). Boxed regions are zoomed in the panels below to highlight the presence (black arrowheads) of interjunctional transerve actin arcs (ITA) only in siCTL cells. Scale bar: 10µm. **B**. Angular distribution of actin filaments in front row siCTL and siPTEN cells (n=300, N=3, Kolmogorov-Smirnov test on the non-normal distribution of data). **C**. Proportion of front row siCTL and siPTEN cells connected by ITA (n=300cells, N=3, paired t-test). **D**. Super resolution immunofluorescence images of actin fibres stained with phalloidin (white) and cell-cell junctions stained with N-cadherin (red) in siCTL and siPTEN migrating astrocytes. Boxes are zoomed-in in the panel below to highlight the shift in orientation of the actin cables in the front row cells. Scale bar:10µm. **E**. Fluorescence images of CCV endothelial cells expressing LifeAct-GFP in CTL and PTENb morphant fish. White arrowheads point at IJAC, in front row migrating cells. Scale bar: 10µm. **F**. Percentage of front row cells from MoCTL and MoPTENb fish connected by IJAC (n=36/29, N=6). paired t-test analysis generated the p-value. **G**. *top:* immunofluorescence image of N-cadherin in migrating astrocyte, where the length (l) and the total contour distance (L) of the lateral cell-cell junctions in a front row cell is drawn to define the linearity index. *bottom:* Junctional linearity index in siCTL and siPTEN front row cells, 8hr after migration.(n=70/65, N=2, unpaired Student t-test). **H**. Junctional linearity index in MoCTL and MoPTENb front row cells, 100min after migration (n=12/16, N=3, unpaired Student t-test). **I**. Scheme summarizing the phenotype seen in front row PTEN depleted cells during collective cell migration. Loss of ITA and less linear junction are highlighted.

To understand the molecular mechanisms responsible for the role of PTEN protein phosphatase activity in the control of actin organisation and collective cell migration, we ran a small protein phosphorylation screen assay. We compared the ratio of protein phosphorylation in siPTEN astrocytes cell vs siCTL with the ratio found in VO-OHpic vs DMSO treated cells, to identify targets of PTEN potentially involved in the control of collective migration. Out of ∼40 proteins of different signalling pathways, AMPKα (T172/T183) phosphorylation was increased by 17% in siPTEN but not in VO-OH treated cells, a difference similar to what is seen for PTEN known target FAK (Y397) (+10%) ^16, 19^ (Supplementary Figure 2A). Western Blot analysis using a different set of AMPKα (T172) phospho-antibody, along with total AMPK measurement, confirmed a strong increase in phosphorylation of AMPKα T172 in PTEN depleted cells (+80%,Figure 3A). Since T172 phosphorylation is known to activate AMPK enzymatic activity ^30^, we analysed the S79 phosphorylation of the biosynthetic kinase acetylCoA carboxylase (ACC), a classic substrate of AMPK^31^. PTEN loss strongly increased ACC phosphorylation, indicating an increased activity of AMPK (Figure 3B). In contrast, cell treatment with LY294002 and VO-OHpic did not affect AMPKα (T172) phosphorylation (Figure 3C) nor its activity (Figure 3D).These data reveal a causal link between PTEN loss and the activation of AMPK, a major guardian of cellular energy levels^32^; and thus more globally a functional link between PTEN and an energy production control hub.

**Figure 3:**
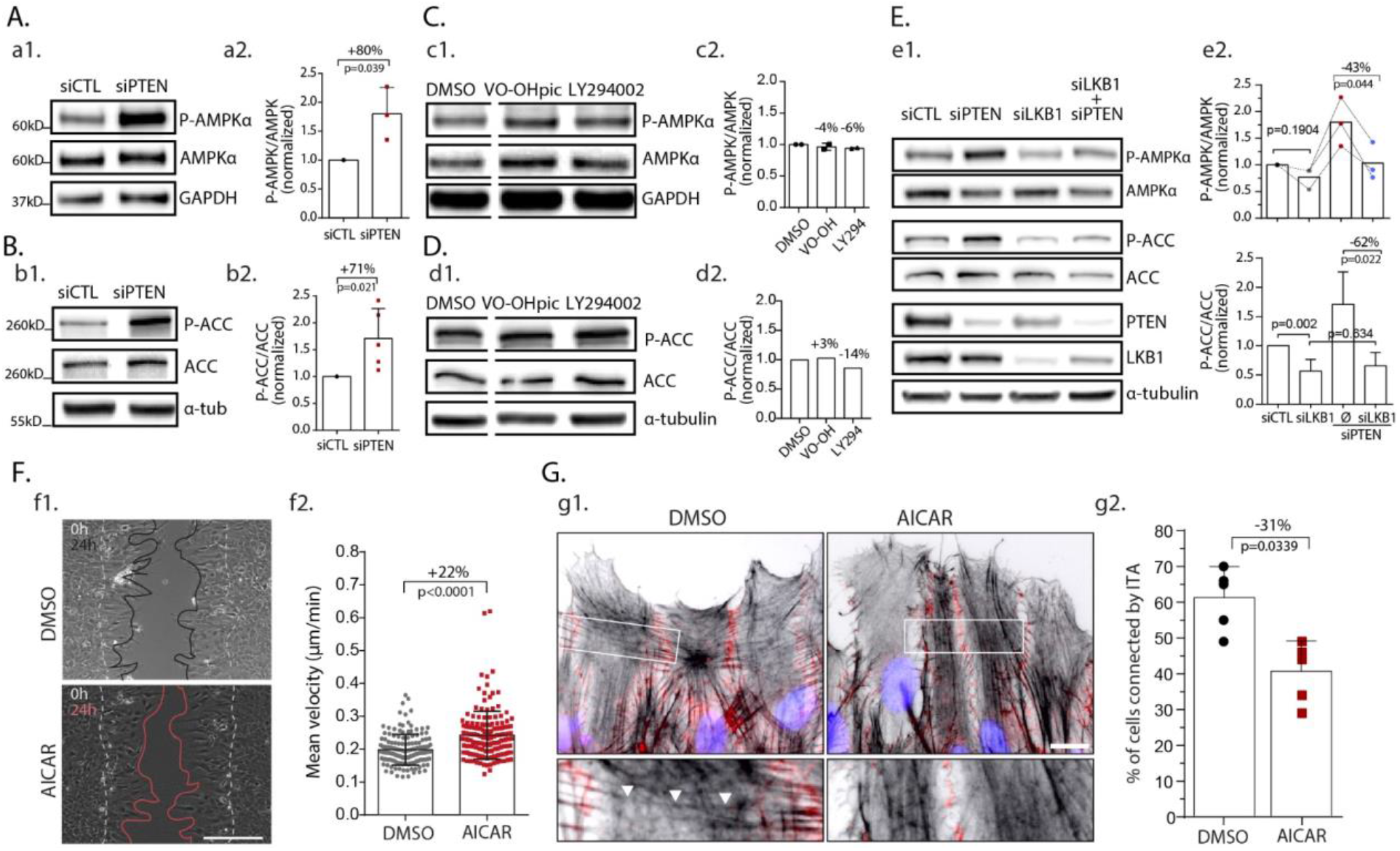
PTEN inhibits AMPK activity to control collective cell migration. **A.C**. Western blot analysis of phosphorylated AMPK (T172), total AMPK, and GAPDH in siCTL and siPTEN (*a1*) and DMSO, VO-OHpic, LY294002-treated (*c1*) astrocytes lysates. *a2*. Normalized ratio (over siCTL) of T172 phosphorylation/total AMPK (+80%, N=2). *c2*. Normalized ratio (over DMSO) of T172 phosphorylation/total AMPK (N=2) **B.D**. Western blot analysis of phosphorylated ACC (S79), total ACC, and α-tubulin in siCTL and siPTEN (*b1*) and DMSO, VO-OHpic, LY294002-treated (*d1*) astrocytes lysates. *b2*. Normalized ratio (over siCTL) of S79 phosphorylation/total ACC (+71%, N=3, paired t-test). *d2*. Normalized ratio (over DMSO) of S79 phosphorylation/total ACC (N=1). **E**. *e1*. Western blot analysis of phosphor-AMPK (T172), total AMPK, ACC, phospho-ACC (S79), PTEN, LKB1 and α-tubulin in siCTL, siPTEN, si LKB1 and siPTEN+siLKB1 astrocytes lysates. *e2*. Normalized ratio (over siCTL) of phospho-AMPK (top) and phospho-ACC (bottom) (N=3, paired t-test). LKB1 depletion rescues basal AMPK activity in siPTEN cells. **F**. *f1*. Phase-contrast images of DMSO and AICAR-treated astrocytes migrating in a wound-healing assay. White dashed lines delineate the border of the wound at t=0. Black/Red lines delineate the border of the monolayer 24h later. Scale bar: 100µm. *f2*. Mean velocity of DMSO and AICAR-treated cells (n=149/157 cells, N=3, unpaired-student t-test). **G**. *g1*. Immunofluorescence images of actin filaments and cell-cell junctions in DMSO and AICAR-treated migrating astrocytes stained with phalloidin (black), N-cadherin (red) and Dapi (blue). Boxed regions are zoomed in the panels below to highlight the presence of ITA only in DMSO cells. Scale bar: 10µm. *g2*. Proportion of front row DMSO and AICAR-treated cells connected by ITA (n=300cells, N=5, paired t-test).

The serine/threonine Liver Kinase B1 (LKB1) genetically and physically interact with PTEN (Supplementary Figure 2B)^33 34^. Interestingly, LKB1 is one of the main kinases phosphorylating AMPK on its T172 residue ^35^. Decreasing LKB1 expression by siRNA in PTEN-depleted astrocytes restored AMPK and ACC phosphorylation to control levels (Figure 3E), showing that LKB1 is involved in the increased AMPK activity induced by PTEN depletion. Of note, LKB1 depletion lowers total PTEN protein level ^36^ without increasing AMPK and ACC phosphorylation (Figure 3E), confirming our conclusion that activation of AMPK in PTEN-depleted cells relies on LKB1. Taken together these data indicate LKB1 acts downstream of PTEN and is responsible for AMPK activation following PTEN loss.

To determine the importance of AMPK overactivation in directing the phenotype of migrating PTEN-depleted cells, we first looked at the effect of AMPK pharmaceutical stimulation on glial cells migrating in a wound-healing assay. Cell treatment with AMPK activator 5-AminoImidazole-4-CArboxamide Ribonucleotide (AICAR), at low dose (40µM), increases cell velocity (+22%) during collective migration (Figure 3F). In these conditions, we also observed a strong decrease in the percentage of cells connected by ITA (Figure 3G), reminiscent of the reorganisation of actin cytoskeleton observed in PTEN-depleted cells.

In search for downstream target of AMPK that could control actin cables and cell-cell junction dynamics, we focused on actin-binding proteins that are present at cell-cell junctions and can be phosphorylated by AMPK. The Vasodilator-Stimulated Phosphoprotein (VASP) met all these criteria. VASP phosphorylation by AMPK occurs on T278 residue *in cellulo* and has been shown to impair actin stress fibres formation in endothelial cells ^37^. AMPK activation following AICAR treatment increased VASP T278 phosphorylation in astrocytes (Figure 4A). In migrating cells, VASP localized both at cell-cell junctions together with actin and N-cadherin and at cell-ECM adhesion sites (Supplementary Figure 3A,B). AMPK activation by AICAR led to VASP delocalization from N-cadherin-mediated adherens junctions (Figure 4B, zoomed-in white boxes) as quantified by a significant drop in the fraction of N-cadherin overlapping VASP at lateral cell-cell contacts (Figure 4C). In addition, we noticed that VASP presence at cell-cell junctions correlated with the presence of ITA (Figure 4B, Supplementary Figure 3C, yellow arrowheads). Interestingly, in AICAR-treated cells, patches of N-cadherin clusters lacking VASP were systematically devoid of ITA (Figure 4B, zoomed-in white boxes).

**Figure 4:**
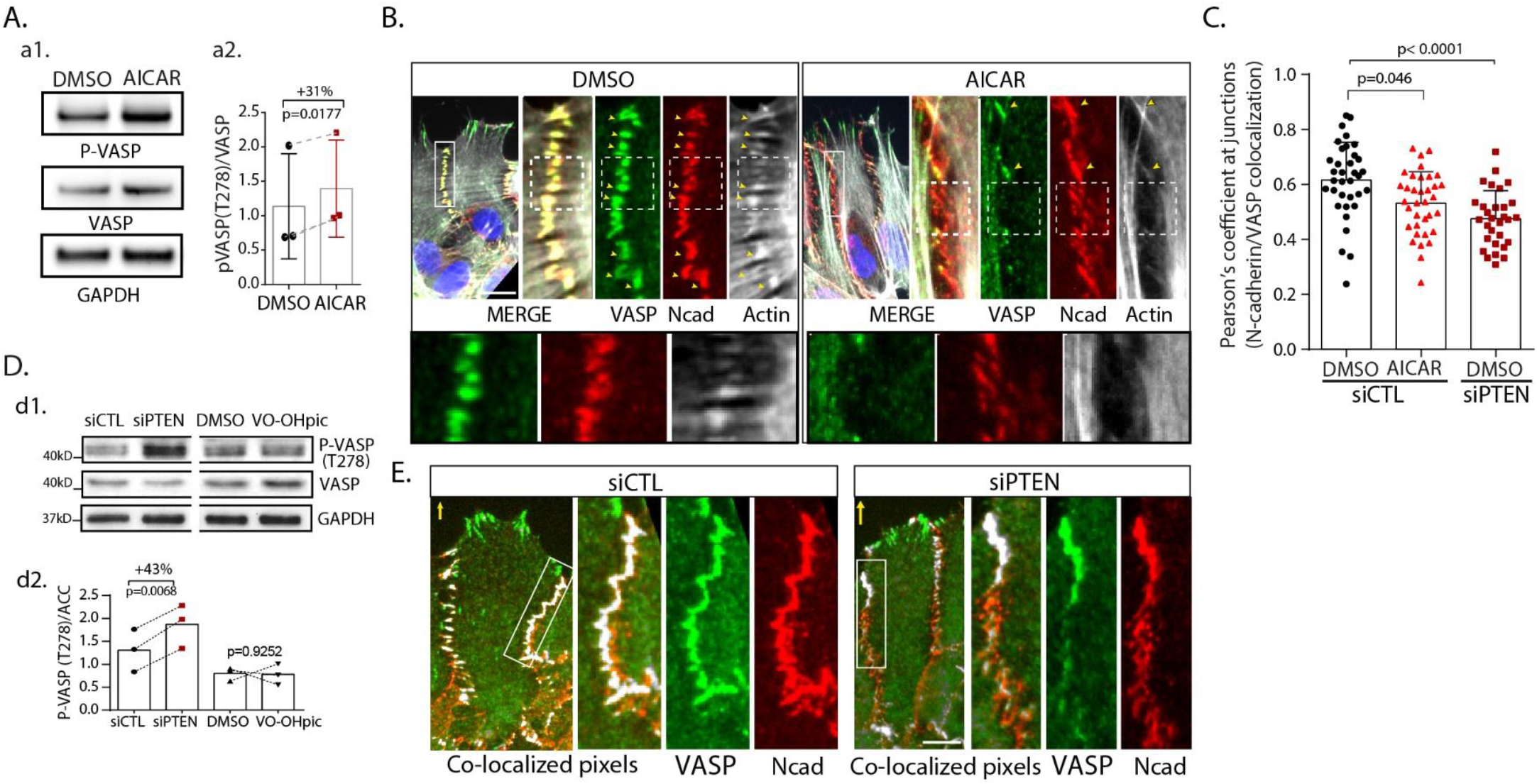
AMPK activation controls VASP phosphorylation and localisation at cell-cell junction. **A**., **D**. *a1*., *d1*. Western blot analysis of phosphorylated VASP (T278), total VASP and GAPDH in DMSO and AICAR treated astrocytes (*a1*) and siCTL, siPTEN, DMSO and VO-OHpic cells (*d1*). *a2*. Ratio of Phospho-VASP over total VASP in DMSO vs AICAR cells (+31%, N=3, paired t-test). *d2*. Ratio of Phospho-VASP over total VASP in siCTL vs siPTEN cells (+43%, N=3, paired t-test) and in DMSO vs VO-OHpic (N=3, paired t-test, no significant difference). **B**. Immunofluorescence images of VASP (green), N-cadherin (red) and F-actin (gray) in DMSO and AICAR treated cells. Cell-cell junctions enriched in VASP (yellow arrowheads) associate with ITA anchoring. White rectangles area are zoomed in the top panel. Dotted-line squares in the zoomed area are further zoomed in the bottom panels. Note that the absence of VASP at cell-cell junctions in AICAR treated cells associates with the absence of ITA. **C**. Fraction of junctional N-cadherin overlapping with VASP in DMSO- and AICAR-treated siCTL cells (n=35, N=3, unpaired t-test) and in DMSO-treated siPTEN cells (n=30, N=3). **E**. Immunofluorescence images of VASP (green), N-cadherin (red) and colocalized pixels between N-cadherin and VASP (gray) in siCTL and siPTEN cells. Scale bars: 10µm.

We then tested whether PTEN depletion, which increases AMPK activity, could affect VASP in a similar way. PTEN loss significantly increased VASP (T278) phosphorylation, contrary to VO-OH-dependent lipid phosphatase inhibition (Figure 4D). PTEN depletion also altered VASP localization at adherens junctions (Figure 4E), but not at focal adhesions (Supplementary Figure 3A,B).We indeed observed a significant drop of the Pearson’s coefficient assessing the colocalization between N-cadherin and VASP in siPTEN cells (Figure 4C). Moreover, similarly to AICAR treated cells, the absence of VASP at cell-cell junctions was systematically associated to the absence of ITA at this specific location (Supplementary Figure 3D, yellow asterisks). Taken together these data reveal that AMPK activation mediates and phenocopies PTEN deletion during collective sheet-like migration. AMPK activation in PTEN-depleted cells increases VASP phosphorylation and triggers its relocalization away from cell-cell junctions, which is associated with destabilisation of interjunctional transverse actin arcs coupling neighbouring leader cells.

Finally, we asked whether AMPK inhibition could inhibit collective migration of PTEN depleted cells. To test this hypothesis, we used siRNA against AMPKα1 to decrease AMPK expression level and activity (Supplementary Figure 4A). Of note, in primary astrocytes, AMPKα depletion does not reduce ACC phosphorylation, indicating that in these cells, AMPK basal activity is low (Figure 5A). However, PTEN depletion increased the basal activity of AMPK as shown by the increased phosphorylation of ACC. In these conditions, depletion of AMPKα1 strongly reduced the pACC/ACC ratio further confirming the role of PTEN in the regulation of AMPK activity (Figure 5A). Depletion of AMPKα1, which did not affect the migration of control astrocytes, led to a significant reduction of wound closure by PTEN-depleted cells (Figure 5B). Cell tracking measurements highlighted that AMPKα depletion reduced the migration speed of PTEN-depleted cells by 24%, which corresponds to a 43% decrease of the increase in cell speed caused by PTEN loss (Figure 5B). Additionally, inhibiting AMPK both with pharmacological inhibitor compound C (CC) or AMPKα1 depletion in PTEN-depleted cells rescued the formation of ITA (Figure 5C) and the colocalization between junctional N-cadherin and VASP (Figure 5D).

**Figure 5:**
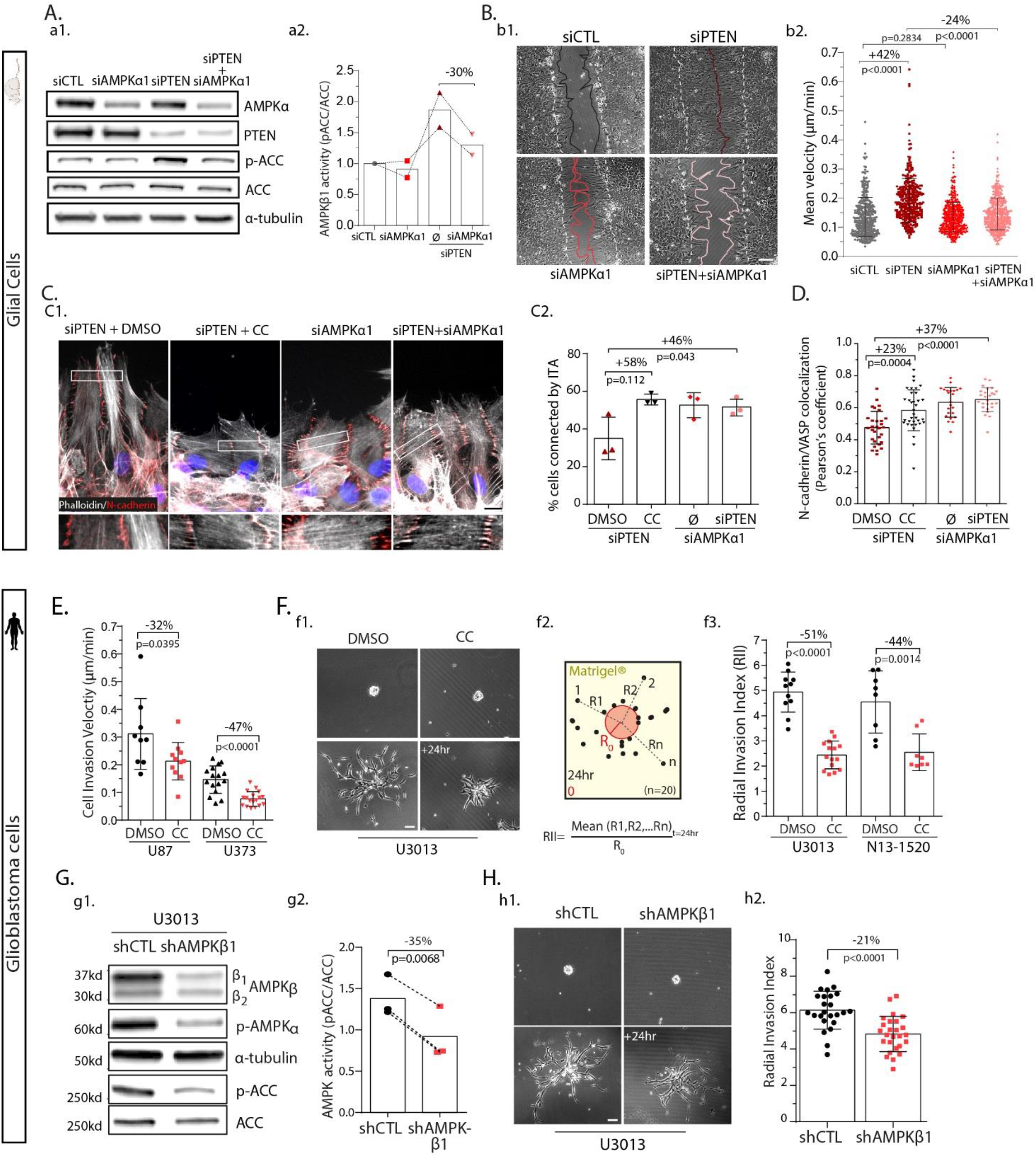
AMPK inhibition reduces collective cell migration and invasion efficiency by restoring leader cell cohesion. **A**. *a1*.Western blot analysis of phosphorylated ACC, total ACC, AMPKα, PTEN and α-tubulin in siCTL, siPTEN, siAMPKα_1_ and siPTEN+siAMPKα_1_ astrocyte lysates. *a2*. Normalized ratio of phosphoACC over total ACC showing efficiency at reducing AMPK activity in double siPTEN+siAMPKα_1_ transfected astrocytes (−30%, N=2). **B**. *b1*. Phase-contrast images of siCTL, siPTEN, siAMPKα_1_ and siPTEN+siAMPKα_1_ astrocyte migrating in a wound-healing assay. White dashed lines delineate the border of the wound at t=0. Coloured lines delineate the border of the monolayer 24h later. Scale bar: 100µm. *b2*. Mean velocity of siCTL, siPTEN, siAMPKα_1_ and siPTEN+siAMPKα_1_ astrocyte during 24hour migration (n=300 cells, N=3, unpaired-student t-test). **C**. *c1*. Immunofluorescence images of actin filaments and cell-cell junctions in siPTEN treated with either DMSO or Compound C (CC), siAMPKα_1_ and siPTEN+siAMPKα_1_ migrating astrocyte stained with phalloidin (gray), N-cadherin (red) and Dapi (blue). Boxed regions are zoomed in the panels below to highlight the rescue of ITA in AMPKα_1_ altered siPTEN cells. Scale bar: 10µm. *c2*. Proportion of leader cells connected by ITA in cells shown in *c1* (n>300, N=3, paired t-test). **D**. Pearson’s coefficient of N-cadherin/VASP colocalisation at lateral cell-cell junctions in cells shown in *c1* (n>25, N=3, unpaired t-test). **E**. Mean cell invasion velocity of U87 and U373 human glioblastoma cell lines treated with or without CC (U87/U373: DMSO, n=9/16; CC, n=11/18; N=3) in a Matrigel® spheroid assay. **F**. *f1*. Phase contrast images of primary GBM cells U3013 just after being included in Matrigel® and 24hr after. Cells are pre-treated with DMSO or CC. Scale bar=100µm. *f2*. Scheme explaining calculation of the radial invasion index (RII) used to measure cell invasion. *f3*. Radial invasion index of U3013 and 1520 primary GBM cells treated with DMSO or CC (U3013/1520: DMSO, n=11/8 spheres; CC, n=15/8 spheres; N=3; unpaired t-test). **G**. *g1*. Western blot analysis of phosphorylated AMPKα, total AMPKβ, phosphorylated ACC, total ACC and α-tubulin on shCTL and shAMPKβ_1_ stable U3013 cells. *g2*. Ratio of phosphoACC/total ACC in shCTL and shAMPKβ_1_ U3013 cells showing reduction of 35% of AMPK activity in shAMPKβ_1_ cells (N=3, paired t-test). **H**. *h1*. Phase contrast images of control primary GBM cells U3013 stably depleted of AMPKβ_1_ (shAMPKβ_1)_ and the control line (shCTL) just after being included in Matrigel® and 24hr after. Scale bar=100µm. *h2*. Radial invasion index of shCTL and shAMPKβ_1_ U3013 cells, showing a 21% decrease in infiltration efficiency when AMPKβ_1_ is depleted (n=24, shCTL, n=26 spheres, shAMPKβ_1_, N=3, unpaired t-test).

The clear inhibitory effect of AMPK inhibition on the migration of PTEN-depleted cells led us to investigate whether inhibition of AMPK could reduce the invasion of PTEN-null cancer cells. Glioblastoma (GBM) are the most common and the most aggressive malignant brain tumours, thought to arise from glial cells at different stages of their differentiation status ^38^. Highly invasive, GBM cells infiltrate the brain collectively in a connected network of cells ^39^ or as diversely cohesive groups or chains of cells along the blood vessels and the myelinated nerve fibres ^40-42^. We first used PTEN null commercial GBM cell lines U87 and U373 ^43^, grown as neurospheres and embedded in Matrigel diluted 1:1. Inhibiting AMPK with Compound C (CC) treatment slowed down invasion speed significantly for both cell lines even though the inhibition appeared more pronounced in U373 (Figure 5E). We then tested the impact of AMPK inhibition in primary patient-derived GBM cells devoid of PTEN (Supplementary Figure 4). AMPK inhibition strongly blocked the radial invasion of U3013 and N13-1520 cells (Figure 5F). To confirm the specific role of AMPK in the blocking of the invasion, we established a stable AMPKβ1-depleted U3013 cell population (shAMPKβ1, Figure 5G, Supplementary Figure 4C). Although AMPK activity was only reduced by 35% (Figure 5G), the invasive capacity of shAMPKβ1 U3013 cells was significantly inhibited (Figure 5H). Together these data show that AMPK is a major actor in controlling migration and invasion in PTEN-depleted cells and suggest that AMPK inactivation may be sufficient to reduce PTEN-null GBM invasion.

## DISCUSSION

In search for oncogenic event that could foster cell motility, we found that loss of tumour suppressor PTEN alone, is sufficient to enhance collective cell migration speed of glial cells *in vitro* and endothelial cells *in vivo*. We provide evidence that it is independent of PI3K/AKT signalling, but requires LKB1-dependent activation of AMPK, a master regulator of metabolism.

Whether collectively migrating PTEN-depleted cells require PI3K/AKT activation to increase their velocity seems to depend both on cell types and on the nature of the migratory stimuli. Contrary to what we observed in PTEN depleted glial cells, and other on PTEN^+/-^ mouse astrocytes^44^, PTEN^-/-^ fibroblasts rely on PI3K/AKT-dependent Rac1 and Cdc42 activation to promote collective motility, in a similar wound-healing assay^11^. In single cell migration, PTEN loss results in PI3K/AKT activation and subsequent Rac1 stimulation in mouse embryonic fibroblasts and neutrophils ^45-47^. The results found in astrocytes may be explained by the presence in these cells of alternative ways of activating Rac1, independent of PI3K/AKT. Alternatively, because cell-cell connective interactions vary between cell types during wound-healing assays, the differences between fibroblasts and astrocytes may rely on front row organisation of the monolayer during wound-healing closure.

Front row CCV endothelial cells and rat astrocytes form tight connections through interjunctionnal transverse actin cables (ITA). We found that front row PTEN-depleted cells lose the ITA-dependent connectivity. It results in less linear junctions possibly due to the decreased intercellular tension or to altered adherens junctions dynamics. The weaker connection between leader cells is associated with increased cell velocity but no alteration of global directionality, presumably due to maintenance of enough cell-cell junction’s integrity. It is in agreement with previous findings showing alteration of ITA affects adherens junction dynamics, which ultimately leads to destabilisation of cell-cell contacts and increased cell velocity^29^.

Mechanistically, we show here that during collective cell migration, PTEN alteration leads to LKB1-dependent activation of AMPK. LKB1 recruits AMPK to E-cadherin rich cell-cell contacts^48, 49^, suggesting here that AMPK activation is spatially restricted at adherens junction in PTEN-depleted cells. VASP is present at adherens junctions in migrating leader cells. Upon PTEN loss, we report that AMPK activation increases VASP phosphorylation on T278, and delocalises it from the adherens junction. In turn, increased T278-phophorylated VASP cytoplasmic accumulation near the adherens junction may alter F-actin elongation^37, 50^, perturbing the formation of the interjunctional actin arcs. Alternatively, AMPK has been shown to regulate actomyosin contractility and junction maturations^48, 51^. Increased contractile forces following AMPK activity may alter local balance of forces at N-cadherin-rich ITA anchoring points and thus participate to their detachment.

Additional pathways controlled by AMPK activation may also contribute to the alteration of collective cell migration. Loss of AMPK was shown to increase surface adhesion and spreading^52^, suggesting increased AMPK activity might have the opposite effect, decreasing matrix attachment and thus promoting migration. Increased FAK (Y397) phosphorylation in PTEN depleted cells would further alter cell-ECM attachment.

PTEN is known to affect cellular bioenergetics and cell growth via its negative regulation of PI3K/AKT-dependent control of mammalian target of rapamycin (mTOR) signalling^53^. We unveil here, in glial cells, a new PI3K-independent function of PTEN in metabolism control, via its up-regulation of AMPK activity. AMPK-dependent metabolic pathways also affects cell migration velocity by regulating intracellular ATP:ADP ratio^54^. AMPK-dependent enhanced energy production at the leading edge of migrating cells also sustains cell motility machinery by controlling polarized trafficking of mitochondria at the front of the cells and lamellipodia turnover^55, 56^.

Our study brings new inputs into how PTEN alteration could drive cancer progression. Because sheet-like migration is often seen in tumour invasive front *in vivo*, notably in skin and intestine tumours ^57^, and in perivascular environment for some GBM cells ^58^, we believe the uncovering of AMPK’s role in mediating the effect of PTEN loss offers a new therapeutic route to tackle cancer cell invasion. While AMPK activity has long been seen as suppressing cancer progression by slowing down cellular growth and proliferation ^59-61^, it was recently shown to be hyperactivated in GBM and promote its development by modifying cellular bioenergetics^62^. We show here that targeting AMPK also reduces GBM cell invasion, reinforcing the interest in developing AMPK inhibitors to treat GBM.

## METHODS

### Zebrafish lines and husbandry

Zebrafish (*Danio Rerio*) of the AB background (Wt, from the Zebrafish International Resource Center) IRC, Eugene, OR, USA), and the transgenic line Tg(fli1a:Lifeact-eGFP)^zf495Tg 63^ were raised according to standard procedures with a 14 hr light/10 hr dark cycle as previously described ^64^. Eggs obtained by natural spawning were bleached and raised at 28°C in Volvic source water supplemented with 280 μg/L of methylene blue (Sigma Aldrich, Cat#: M4159). N-Phenylthiourea (PTU, Sigma Aldrich, Cat#: P7629) was added to raising medium (0.003% final) from 24 hpf onwards to prevent pigmentation and facilitate imaging. Animal experiments were conducted according to European Union guidelines for handling of laboratory animals (http://ec.europa.eu/environment/chemicals/lab_animals/home_en.htm). All protocols were approved by the Ethical Committee for Animal Experimentation of Institut Pasteur – CEEA 89 and the French Ministry of Research and Education (permit #01265.03). During injections or live imaging sessions, animals were anaesthetized with Tricaine (Ethyl 3-aminobenzoate methanesulfonate, Sigma-Aldrich, Cat#: A5040). At the end of the experimental procedures, they were euthanized by anaesthetic overdose.

### Mammalian cell culture

Primary astrocytes were obtained from E17 rat embryos as previously described^20^. Cells were grown in 1g/L D-glucose DMEM+Pyruvate supplemented with 10% FBS (ThermoFischer Scientific, Waltham, MA), penicillin-streptomycin (100 U.ml^−1^ and 100 μg.ml^−^1, Gibco™) and Amphotericin B (2.5µg/ml, Gibco™) at 5% CO2 and 37°C. Medium was changed 1 day after transfection and 1 day before the experiments. Human Neural Stem Cells (hNSC) were obtained from ThermoFischer Scientific (Gibco™, ref#N7800100, H9-derived) and cultured as recommended using KnockOut™ DMEM/F-12 Basal Medium, StemPro™ Neural Supplement, human FGF-basic and human EGF (animal free, Peprotech) at 20 ng/mL. Commercial glioblastoma cell lines U87 and U373 were grown in modified Eagle’s medium (MEM) supplemented with 10% FBS, penicillin–streptomycin and non-essential amino-acids (all from Gibco™). U3013 and N13-1520 primary human glioblastoma cells were acquired respectively from The Human Glioblastoma Cell Culture Resource (Uppsala University, Sweden, www.hgcc.se)^65^ and GlioTEx (Institut du Cerveau et de la Moelle Epinière, ICM, F-75013, Paris, France)^66^; both institutions having the necessary ethical agreements to collect GBM samples from informed patients. Once thawed, both cell types were plated on Geltrex™ pre-coated plates (Gibco) and grown using serum-free GBM complete medium consisting in Neurobasal-A medium mixed 1:1 with DMEM/F12+GlutaMAX and complemetend with human FGF-basic and human EGF (animal free, Peprotech) at 20 ng/mL, and B-27® (Gibco). HEK-293T were grown using DMEM + GlutaMAX + 4.5g/L D-glucose + Pyruvate (Gibco™) medium complemented with 10% FBS and penicillin-streptomycin. Cells were routinely tested for mycoplasma contamination using PCR detection kits.

### Gene depletion and transfection protocols

Astrocytes were transfected with siRNAs (1-5nM) or plasmids (5µg) using a Nucleofector machine (Lonza) and the appropriate Lonza glial cell nucleofector solution. Transfected cells were then plated on appropriate supports previously coated with poly-L-Ornythin (Sigma) and experiments were performed 4 days post-transfection, when optimal protein silencing or expression was observed. Sequences of siRNAs used here are: siCTL (luciferase): UAAGGCUAUGAAGAGAUAC; siPTEN: AGGACGGACUGGUGUAAUGUU; siLKB1: GCUCUUUGAGAACAUCGGG. siAMPKα1 consisted in the ON-TARGETplus SMARTpool against rat PRKAA1 (Dharmacon™,Ref#SO-2905147G). PTEN rescue experiments were realised using PTENwt and PTENc124s cloned into pGK-GFP plasmids, from original pCGN-HA-PTENwt/c124s plasmids from N.Tonks. pEGFP-C1-LKB1 (LKB1-GFP) plasmid was kindky provided by M.Salmi. N-cadherin retrograde flow was measured after transfection with pEGFP-C1-N-cadherin (Ncad-GFP)^29^. To generate stable primary human GBM cells devoid of AMPK (shAMPKβ1) and the shCTL control clone, GBM#U3013 cells were infected with lentiviral particles generated by transfecting HEK293 cells with pLKO.1-puro plasmids from the Mission shRNA library (Sigma-Aldrich). Briefly, lentiviral particles were added to the plated cells for 24h, before cells were washed with GBM complete medium. 2days later 3µg/ml puromycin was added to select positively infected cells. Antibiotic selection was prolonged for several days until the separate uninfected plated cells, seeded at the same concentration, were all dead. Several shRNA sequences against AMPKβ1 from the Mission library were tested and the one inducing the maximal protein depletion was kept (Ref#TRCN0000004770). pLKO.1-puro non-target shRNA (Sigma-Aldrich) was used to produce the shCTL clone.

### Lentivirus production

Lentiviral particles were generated by transient transfection of HEK 293T cells. In brief, 7× 10^6^ cells were transfected with a mixture of DNA/CaCl_2_ diluted 1:1 in 20 mmol.l^−1^ HEPES after 20min of incubation at room temperature. DNA comprised a mix of 10µg 2^nd^ generation packaging plasmid psPAX2, 5µg of viral envelope plasmid pMD2.G and 10µg of plasmid of interest (pLKO.1-puro or pGK-PTEN plasmids). Viral supernatant was harvested 36 and 48 h post-transfection and concentrated using ultracentrifugation (Beckman Coulter, Optima XPN-80) for 1.5 h at 19 000 rpm and 4 °C and then stored at −80o C. Viral particles were then dropped directly in the targeted cells medium, at 1 in 5000 dilution.

### Zebrafish morpholino microinjections

Zebrafish embryos were injected using a Picospritzer III microinjector (Parker Hannifin) and a mechanical micromanipulator (M-152; Narishige, Tokyo, Japan). Morpholinos were loaded in a pulled borosilicate glass filament containing capillary (GC100F-15; Harvard Apparatus, Holliston). Injections of typically 1nL were performed at 1 cell stage embryos, at room temperature. The sequences of the Morpholinos (Gene Tools, LLC, Philomath, OR, USA) are: PTENa: CCTCGCTCACCCTTGACTGTGTATG; PTENb: CTTTCGGACGGTCGGTCGTCTTTA. A Standard Control oligo from the company was used as control.

### Drugs Treatments and chemicals

Drugs were added one hour before the start of the experiments. VO-OHpic (Tebu-bio, Ref#B-0351), Compound C (CC) also called Dorsomorphin (EMD, Millipore, Ref#171261), and LY294002 (Calbiochem™) were used at 10µM. AICAR (Enzo, Ref#BML-EI330-0050) was used at a low dose of 40µM.

### *In vitro* Migration and Invasion Assays

For scratch-induced wound-healing migration assays, cells were seeded on poly-L-ornithine coated coverslips (for immunofluorescence), 35-mm-diameter glass-bottom culture dishes (MatTek®) (for fluorescent videomicroscopy) or 12-well plates (for brightfield videomicroscopy), and grown to confluence. On the day of the experiment, the monolayer of cells is scratched with a blunt-ended microinjection needle, creating a 300/500nm-wide wound that is closed up by cell’s collective migration. For immunofluorescence staining, cells are allowed to migrate for 8hours. For fluorescent videomicroscopy and N-cadherin flow measurement, cells are let to migrate for 4 hours prior to imaging to allow extension of their protrusion. Typically, two hours-long movies are recorded to analyse N-cadherin dynamics using a spinning-disk confocal microscope (UltraVIEW vox, PerkinElmer) composed of a Zeiss AxioObserver Z1 stand equipped with a spinning-disk head Yokagawa CSUX1, two EMCCD cameras (Hamamatsu, Japan) and a humidified, C0_2_ controlled, heating chamber. The microscope is monitored by the Volocity® 3D image analysis software (PerkinElmer). To assess collective cell migration kinetics, 24hours movies with a 15min timelapse interval are recorded using brightfield videomicroscopy performed by a Nikon Eclipse Ti2 epifluorescence inverted microscope equipped with a pco.edge 3.1 sCMOS camera (PCO, Germany) in a humidified, CO_2_-controlled heating chamber (5% CO2 and 37°C, Okolab, Italy). All images were acquired with a dry 10x 0.45 NA objective by the MetaMorph® Microscopy Automation and Image Analysis Software (Molecular Devices, CA, USA).

Radial 3D invasion assays were performed by embedding glioblastoma spheroids into a 50% Matrigel® solution (Corning®, Merck) (1:1 Matrigel® diluted in spheroids + GBM medium). Their efficiency at disseminating within the gel is analysed for 24 hours, by acquiring brightfield images every 15min. Glioblastoma spheroids are generated by growing GBM cells in non-adherent flasks with the same glioblastoma complete medium for a minimum of 2-3 days, until the spheroids reach ∼100/200µm in diameter.

### *In vivo* Migration

For *in vivo* imaging, five to ten 24hour-post-fecundation zebrafish embryos were manually dechorionated with forceps, anaesthetized with 112 µg/ml Tricaine, immobilized in 1% low-melting-point agarose supplemented with 1xTricaine, in the centre of a 35-mm glass-bottomed dishes (MatTek Life Sciences, MA, USA), and then covered with ∼2 ml Volvic water containing 0.2x Tricaine. Fluorescence imaging of the Tg(Lifeact::gfp) strain was performed using the UltraVIEW vox microscope with either a 63× or a 40x oil-immersion objective to collect 0.3µm z-stack images every 2 minutes using the Volocity® software.

### Migration and Invasion Kinetics Measurement

Manual Tracking plugin (FIJI, ImageJ ^67^) was used to monitor collective cell migration and 3D Matrigel® invasion characteristics by tracking the nucleus of non-dividing leader cells located at the wound/spheroid edge. Velocity, directionality and persistence of direction were calculated following a previously published protocol^68^. One hundred randomly chosen cells were analysed per experiment. When differences in cell invasion efficiency were obvious, a radial invasion index was calculated based on t0 and t+24h images, as explained in the Figure4. Briefly, the mean radius (i.e, the distance between the cell nucleus region to the centre of the spheroid) at 24hours of the 20 most spread cells was measured and normalized by the radius of the spheroid at t0.

### Immunofluorescence

Cells migrating for 8h were fixed in cold methanol for 3 min at −20^°^C or 4% PFA for 10min at 37^°^C, permeabilised for 10 min with 0.2% Triton and blocked with 3% BSA in PBS for 1h at room temperature. Cells were then incubated for 1h with primary antibodies diluted in PBS 1X, washed three times in PBS, and incubated another hour with secondary antibodies. Finally, coverslips were washed and mounted in ProLong Gold with DAPI (Thermo Fisher Scientific). The following primary antibodies were used: anti–α-tubulin (MCA77G YL1/2, rat monoclonal; Bio-Rad), anti-GAPDH (mouse, Chemicon International, MAB374), anti-Paxillin (mouse, BD transduction, #610051), anti–N-cadherin (ab12221, rabbit polyclonal, lot GR139340-26, and sc-31030, goat polyclonal, clone K-20, lot B2014; Abcam), anti-P-Akt (Rabbit, Cell Signaling 4060S), anti-Akt (rabbit, Cell Signaling 4685S), anti-AMPKα (Rabbit, Cell Signaling 2532S), anti-AMPKβ (Rabbit, Cell Signaling 4150S), P-AMPKα/T172 (Rabbit, Cell Signaling 2585S), ACC (Rabbit, Cell Signaling, 3662S), P-ACC (Rabbit, Cell Signaling 3661S), LKB1 (Rabbit, Cell Signaling, 3047),VASP (Rabbit, Cell Signaling 3132), P-VASP/T278 (Rabbit, Sigma Aldrich, SAB4200521), GFP-HRP (Abcam, Ab6663). Phalloidin-iFluor647 reagent (Abcam Ab176759) was used to label F-actin filaments. Secondary antibodies were Alexa Fluor 488 donkey anti–rabbit (711-545-152), Rhodamine (TRITC) donkey anti–rabbit (711-025-152), Rhodamine (TRITC) donkey anti-mouse (715-025-151), Alexa Fluor 647 donkey anti-rabbit (711-695-152), Alexa Fluor 647 donkey anti–goat (705-605-147), and Alexa Fluor 488 donkey anti–rat (712-545-153); all from Jackson ImmunoResearch. Epifluorescence images were obtained on a microscope (DM6000; Leica Biosystems) equipped with 40×, 1.25 NA, and 63×, 1.4 NA, objective lenses and were recorded on a charge-coupled device camera using Leica Biosystems software. Super-resolution images (Figure 2D) were acquired xith Zeiss LSM780 ELYRA with 63X 1.4 NA or 100X 1.46 NA objectives and recorded on an EMCCD Andor Ixon 887 1K with Zen software.

### Electrophoresis and Western blot

Cells are lysed with Laemmli buffer composed of 60mM Tris pH6.8, 10% glycerol, 2% SDS and 50mM DTT with the addition of either a 10x phosphatase cocktail inhibitor made of 200mM Napp, 500mM NaF, 10mM sodium orthovanadate and 20mM EDTA, or PhosSTOP™ (Sigma-Aldrich). Samples are then boiled for 5 minutes at 95^°^C before loading on polyacrylamide gels. Transfer is performed at 0.1A constant overnight or at 0.3A for 2h at 4^°^C on nitrocellulose membranes. Finally, membranes are blocked with 5% milk or BSA for phosphorylated proteins in TBS+0.2% Tween®20 (Sigma-Aldrich) for 1h and incubated 1h with primary antibody at room temperature. After being washed 3 times in TBST, they are incubated 1h with HRP-conjugated secondary antibody. Bands are revealed with ECL chemoluminescence substrate (Pierce, Thermoscientific).

### Immunoprecipitation

Confluent 10cmØ dishes of transfected HEK293 cells s were washed with cold 1xPBS and lysed with 1ml of 1x IP buffer (500 mM Tris HCL pH 7.5, Triton 20%, 2M NaCl) with the fresh addition of cOmplete™ protease inhibitor cocktail (Roche). Lysates were centrifuged at 13.000 rpm 2.30 min at 4^°^C. Some supernatant mixed 1:1 with 2x Laemmli buffer was stored at −20^°^C to serve as Input loading control. The rest of the supernatant was incubated for 2h at 4^°^C on the spinning wheel with Protein G beads (50ul/dish) and 1µg of homemade GFP-GST nanobodies collected from BL21 bacteria transfected with pGEX-GST-GFP, or GST beads only. Beads were then washed eight times with IP washing buffer (50mM Tris HCL pH 7.5, 150mM NaCl, 1mM EDTA, 2.5mM MgCl2) before adding Laemmli buffer and loading on precast gels (Invitrogen).

### Proteome profiler array

The human Phospho-kinase array kit (R&D Systems, Ref#ARY003B) was used to perform the small phosphoprotemics screen to unveil new targets affected by PTEN depletion. Experiments and analysis were realized in accordance with the provider’s protocol.

### Immunofluorescence Image analysis

*Angular distribution of actin filaments* in cells was measured as follow. After defining a reference orientation parallel to the wound, leader cells were segmented manually based on the N-cadherin staining and OrientationJ plugin was used to extract local orientations of F-actin filaments based on Phalloidin staining. Kolmogorov-Smirnov test on the non-normal distribution of data was performed to validate the differences between siCTL and siPTEN.

#### Interjunctional actin arcs (IJACs)

Leader cells in the front row were scored manually as being connected by IJACs if at least two actin arcs could see anchored at cell-cell junctions on both side of the cell.

#### VASP/Ncad colocalization

VASP presence at cell-cell junctions was monitored by measuring its colocalisation with junctional N-cadherin. ROI was drawn around lateral cell-cell border from leading edge to roughly just in front of the nucleus. Then, the Paersons Coefficient and the Overlapping Coefficient (fraction of protein 1 overlapping with protein 2) were measured on thresholded immunofluorescence images using JACOP (Just Another Colocalization Plugin) plugin in FIJI. One data point corresponds to the mean of the left/right junction.

*Lateral cell-cell junction linearity* was defined as the ratio between the length (straight line between the further at the rear to the most in front cell-cell contact) and the actual distance of the lateral intercellular contacts (cell-cell junction contour), based on the N-cadherin staining. Linearity index is 1 when the cell-cell junction is perfectly straight.

All data are presented as the mean +/- standard deviation of at least three independent experiments.

### Statistical analyses

Statistical analysis were obtained with two-tailed unpaired or paired Student’s t-test depending on the type of experiment conducted or with one-way ANOVA (Analysis Of Variance), when data followed a Gaussian distribution. Otherwise a Mann-Whittney non parametric analysis was performed. Quantification and statistical analysis were realised using GraphPad Prism software and p-values measurement from the appropriate statistical tests appear on each graph. Error bars on each graph represent standard deviation.

## ACKNOWLEDGMENT

We are extremely grateful to Dr P.Herbomel (Institut Pasteur, France) and his lab for providing us with the zebrafish lines and for their valuable expertise on zebrafish experimental procedures. We also would like to thank the Pasteur Imaging plateform (Imagopole, C2RT) and Dmitri Ershov from the Biostatistics and Bioinformatic Hub (Institut Pasteur) and Image Analysis Hub (Institut Pasteur, C2RT), for help with the image and data analysis. We thank Dr Cécile Leduc for her help with super-resolution imaging. imaging. We thank the Etienne-Manneville’s lab for helpful input on the manuscript. F.P is financially supported by the Fondation TOURRE and the Fondation ARC pour la Recherche contre le Cancer. L.C

## AUTHOR CONTRIBUTIONS

F.P, L.C, S.E.M conceived the experiments, and together with I.P, B.B and F.L carried it out; F.P, L.C and B.B designed and carried out the data analysis; F.P. and S.E.M co-wrote the paper. K.F.N and ICM provided the primary glioblastoma cells.

## REFERENCES

1. Hollander, M.C., Blumenthal, G.M. & Dennis, P.A. PTEN loss in the continuum of common cancers, rare syndromes and mouse models. Nat Rev Cancer 11, 289–301 (2011).

2. Lee, Y.R., Chen, M. & Pandolfi, P.P. The functions and regulation of the PTEN tumour suppressor: new modes and prospects. Nat Rev Mol Cell Biol 19, 547–562 (2018).

3. Cancer Genome Atlas Research, N. Comprehensive genomic characterization defines human glioblastoma genes and core pathways. Nature 455, 1061–1068 (2008).

4. Verhaak, R.G. et al. Integrated genomic analysis identifies clinically relevant subtypes of glioblastoma characterized by abnormalities in PDGFRA, IDH1, EGFR, and NF1. Cancer Cell 17, 98–110 (2010).

5. Brennan, C.W. et al. The somatic genomic landscape of glioblastoma. Cell 155, 462–477 (2013).

6. Maehama, T. & Dixon, J.E. The tumor suppressor, PTEN/MMAC1, dephosphorylates the lipid second messenger, phosphatidylinositol 3,4,5-trisphosphate. J Biol Chem 273, 13375–13378 (1998).

7. Myers, M.P. et al. P-TEN, the tumor suppressor from human chromosome 10q23, is a dual-specificity phosphatase. Proc Natl Acad Sci U S A 94, 9052–9057 (1997).

8. Song, M.S., Salmena, L. & Pandolfi, P.P. The functions and regulation of the PTEN tumour suppressor. Nat Rev Mol Cell Biol 13, 283–296 (2012).

9. Stambolic, V. et al. Negative regulation of PKB/Akt-dependent cell survival by the tumor suppressor PTEN. Cell 95, 29–39 (1998).

10. Tu, T., Chen, J., Chen, L. & Stiles, B.L. Dual-Specific Protein and Lipid Phosphatase PTEN and Its Biological Functions. Cold Spring Harb Perspect Med 10 (2020).

11. Liliental, J. et al. Genetic deletion of the Pten tumor suppressor gene promotes cell motility by activation of Rac1 and Cdc42 GTPases. Curr Biol 10, 401–404 (2000).

12. Funamoto, S., Meili, R., Lee, S., Parry, L. & Firtel, R.A. Spatial and temporal regulation of 3-phosphoinositides by PI 3-kinase and PTEN mediates chemotaxis. Cell 109, 611–623 (2002).

13. Iijima, M. & Devreotes, P. Tumor suppressor PTEN mediates sensing of chemoattractant gradients. Cell 109, 599–610 (2002).

14. Friedl, P. & Gilmour, D. Collective cell migration in morphogenesis, regeneration and cancer. Nat Rev Mol Cell Biol 10, 445–457 (2009).

15. Leslie, N.R., Yang, X., Downes, C.P. & Weijer, C.J. PtdIns(3,4,5)P(3)-dependent and -independent roles for PTEN in the control of cell migration. Curr Biol 17, 115–125 (2007).

16. Tamura, M. et al. Inhibition of cell migration, spreading, and focal adhesions by tumor suppressor PTEN. Science 280, 1614–1617 (1998).

17. Raftopoulou, M., Etienne-Manneville, S., Self, A., Nicholls, S. & Hall, A. Regulation of cell migration by the C2 domain of the tumor suppressor PTEN. Science 303, 1179–1181 (2004).

18. Gildea, J.J. et al. PTEN can inhibit in vitro organotypic and in vivo orthotopic invasion of human bladder cancer cells even in the absence of its lipid phosphatase activity. Oncogene 23, 6788–6797 (2004).

19. Tamura, M., Gu, J., Takino, T. & Yamada, K.M. Tumor suppressor PTEN inhibition of cell invasion, migration, and growth: differential involvement of focal adhesion kinase and p130Cas. Cancer Res 59, 442–449 (1999).

20. Etienne-Manneville, S. In vitro assay of primary astrocyte migration as a tool to study Rho GTPase function in cell polarization. Methods Enzymol 406, 565–578 (2006).

21. Helker, C.S. et al. The zebrafish common cardinal veins develop by a novel mechanism: lumen ensheathment. Development 140, 2776–2786 (2013).

22. Croushore, J.A. et al. Ptena and ptenb genes play distinct roles in zebrafish embryogenesis. Dev Dyn 234, 911–921 (2005).

23. Faucherre, A., Taylor, G.S., Overvoorde, J., Dixon, J.E. & Hertog, J. Zebrafish pten genes have overlapping and non-redundant functions in tumorigenesis and embryonic development. Oncogene 27, 1079–1086 (2008).

24. Mak, L.H., Vilar, R. & Woscholski, R. Characterisation of the PTEN inhibitor VO-OHpic. J Chem Biol 3, 157–163 (2010).

25. Rosivatz, E. et al. A small molecule inhibitor for phosphatase and tensin homologue deleted on chromosome 10 (PTEN). ACS Chem Biol 1, 780–790 (2006).

26. Mayor, R. & Etienne-Manneville, S. The front and rear of collective cell migration. Nat Rev Mol Cell Biol 17, 97–109 (2016).

27. Rorth, P. Fellow travellers: emergent properties of collective cell migration. EMBO Rep 13, 984–991 (2012).

28. Mishra, A.K., Campanale, J.P., Mondo, J.A. & Montell, D.J. Cell interactions in collective cell migration. Development 146 (2019).

29. Peglion, F., Llense, F. & Etienne-Manneville, S. Adherens junction treadmilling during collective migration. Nat Cell Biol 16, 639–651 (2014).

30. Hong, S.P., Leiper, F.C., Woods, A., Carling, D. & Carlson, M. Activation of yeast Snf1 and mammalian AMP-activated protein kinase by upstream kinases. Proc Natl Acad Sci U S A 100, 8839–8843 (2003).

31. Munday, M.R., Campbell, D.G., Carling, D. & Hardie, D.G. Identification by amino acid sequencing of three major regulatory phosphorylation sites on rat acetyl-CoA carboxylase. Eur J Biochem 175, 331–338 (1988).

32. Hardie, D.G. AMP-activated/SNF1 protein kinases: conserved guardians of cellular energy. Nat Rev Mol Cell Biol 8, 774–785 (2007).

33. Stratakis, C.A. et al. Carney complex, Peutz-Jeghers syndrome, Cowden disease, and Bannayan-Zonana syndrome share cutaneous and endocrine manifestations, but not genetic loci. J Clin Endocrinol Metab 83, 2972–2976 (1998).

34. Mehenni, H. et al. LKB1 interacts with and phosphorylates PTEN: a functional link between two proteins involved in cancer predisposing syndromes. Hum Mol Genet 14, 2209–2219 (2005).

35. Woods, A. et al. LKB1 is the upstream kinase in the AMP-activated protein kinase cascade. Curr Biol 13, 2004–2008 (2003).

36. Jimenez, A.I., Fernandez, P., Dominguez, O., Dopazo, A. & Sanchez-Cespedes, M. Growth and molecular profile of lung cancer cells expressing ectopic LKB1: down-regulation of the phosphatidylinositol 3’-phosphate kinase/PTEN pathway. Cancer Res 63, 1382–1388 (2003).

37. Blume, C. et al. AMP-activated protein kinase impairs endothelial actin cytoskeleton assembly by phosphorylating vasodilator-stimulated phosphoprotein. J Biol Chem 282, 4601–4612 (2007).

38. Ostrom, Q.T. et al. CBTRUS statistical report: Primary brain and central nervous system tumors diagnosed in the United States in 2006-2010. Neuro Oncol 15 Suppl 2, ii1–56 (2013).

39. Osswald, M. et al. Brain tumour cells interconnect to a functional and resistant network. Nature 528, 93–98 (2015).

40. Cuddapah, V.A., Robel, S., Watkins, S. & Sontheimer, H. A neurocentric perspective on glioma invasion. Nat Rev Neurosci 15, 455–465 (2014).

41. Seano, G. & Jain, R.K. Vessel co-option in glioblastoma: emerging insights and opportunities. Angiogenesis 23, 9–16 (2020).

42. Hong, J.H. et al. Modulation of Nogo receptor 1 expression orchestrates myelin-associated infiltration of glioblastoma. Brain 144, 636–654 (2021).

43. Li, Y. et al. PTEN has tumor-promoting properties in the setting of gain-of-function p53 mutations. Cancer Res 68, 1723–1731 (2008).

44. Dey, N. et al. The protein phosphatase activity of PTEN regulates SRC family kinases and controls glioma migration. Cancer Res 68, 1862–1871 (2008).

45. Kolsch, V., Charest, P.G. & Firtel, R.A. The regulation of cell motility and chemotaxis by phospholipid signaling. J Cell Sci 121, 551–559 (2008).

46. Higuchi, M., Masuyama, N., Fukui, Y., Suzuki, A. & Gotoh, Y. Akt mediates Rac/Cdc42-regulated cell motility in growth factor-stimulated cells and in invasive PTEN knockout cells. Curr Biol 11, 1958–1962 (2001).

47. Subramanian, K.K. et al. Tumor suppressor PTEN is a physiologic suppressor of chemoattractant-mediated neutrophil functions. Blood 109, 4028–4037 (2007).

48. Bays, J.L., Campbell, H.K., Heidema, C., Sebbagh, M. & DeMali, K.A. Linking E-cadherin mechanotransduction to cell metabolism through force-mediated activation of AMPK. Nat Cell Biol 19, 724–731 (2017).

49. Sebbagh, M., Santoni, M.J., Hall, B., Borg, J.P. & Schwartz, M.A. Regulation of LKB1/STRAD localization and function by E-cadherin. Curr Biol 19, 37–42 (2009).

50. Benz, P.M. et al. Differential VASP phosphorylation controls remodeling of the actin cytoskeleton. J Cell Sci 122, 3954–3965 (2009).

51. Lee, J.H. et al. Energy-dependent regulation of cell structure by AMP-activated protein kinase. Nature 447, 1017–1020 (2007).

52. Georgiadou, M. et al. AMPK negatively regulates tensin-dependent integrin activity. J Cell Biol 216, 1107–1121 (2017).

53. Aquila, S. et al. The Tumor Suppressor PTEN as Molecular Switch Node Regulating Cell Metabolism and Autophagy: Implications in Immune System and Tumor Microenvironment. Cells 9 (2020).

54. Zanotelli, M.R. et al. Regulation of ATP utilization during metastatic cell migration by collagen architecture. Mol Biol Cell 29, 1–9 (2018).

55. Cunniff, B., McKenzie, A.J., Heintz, N.H. & Howe, A.K. AMPK activity regulates trafficking of mitochondria to the leading edge during cell migration and matrix invasion. Mol Biol Cell 27, 2662–2674 (2016).

56. Schuler, M.H. et al. Miro1-mediated mitochondrial positioning shapes intracellular energy gradients required for cell migration. Mol Biol Cell 28, 2159–2169 (2017).

57. Clark, A.G. & Vignjevic, D.M. Modes of cancer cell invasion and the role of the microenvironment. Curr Opin Cell Biol 36, 13–22 (2015).

58. Gritsenko, P., Leenders, W. & Friedl, P. Recapitulating in vivo-like plasticity of glioma cell invasion along blood vessels and in astrocyte-rich stroma. Histochem Cell Biol 148, 395–406 (2017).

59. Vara-Ciruelos, D., Russell, F.M. & Hardie, D.G. The strange case of AMPK and cancer: Dr Jekyll or Mr Hyde? (dagger). Open Biol 9, 190099 (2019).

60. Inoki, K., Zhu, T. & Guan, K.L. TSC2 mediates cellular energy response to control cell growth and survival. Cell 115, 577–590 (2003).

61. Gwinn, D.M. et al. AMPK phosphorylation of raptor mediates a metabolic checkpoint. Mol Cell 30, 214–226 (2008).

62. Chhipa, R.R. et al. AMP kinase promotes glioblastoma bioenergetics and tumour growth. Nat Cell Biol 20, 823–835 (2018).

63. Phng, L.K., Stanchi, F. & Gerhardt, H. Filopodia are dispensable for endothelial tip cell guidance. Development 140, 4031–4040 (2013).

64. Kimmel, C.B., Ballard, W.W., Kimmel, S.R., Ullmann, B. & Schilling, T.F. Stages of embryonic development of the zebrafish. Dev Dyn 203, 253–310 (1995).

65. Xie, Y. et al. The Human Glioblastoma Cell Culture Resource: Validated Cell Models Representing All Molecular Subtypes. EBioMedicine 2, 1351–1363 (2015).

66. Rosenberg, S. et al. Multi-omics analysis of primary glioblastoma cell lines shows recapitulation of pivotal molecular features of parental tumors. Neuro Oncol 19, 219–228 (2017).

67. Schindelin, J. et al. Fiji: an open-source platform for biological-image analysis. Nat Methods 9, 676–682 (2012).

68. De Pascalis, C. et al. Intermediate filaments control collective migration by restricting traction forces and sustaining cell-cell contacts. J Cell Biol 217, 3031–3044 (2018).

