## Supplementary Information for "PTEN inhibits AMPK to control collective migration"

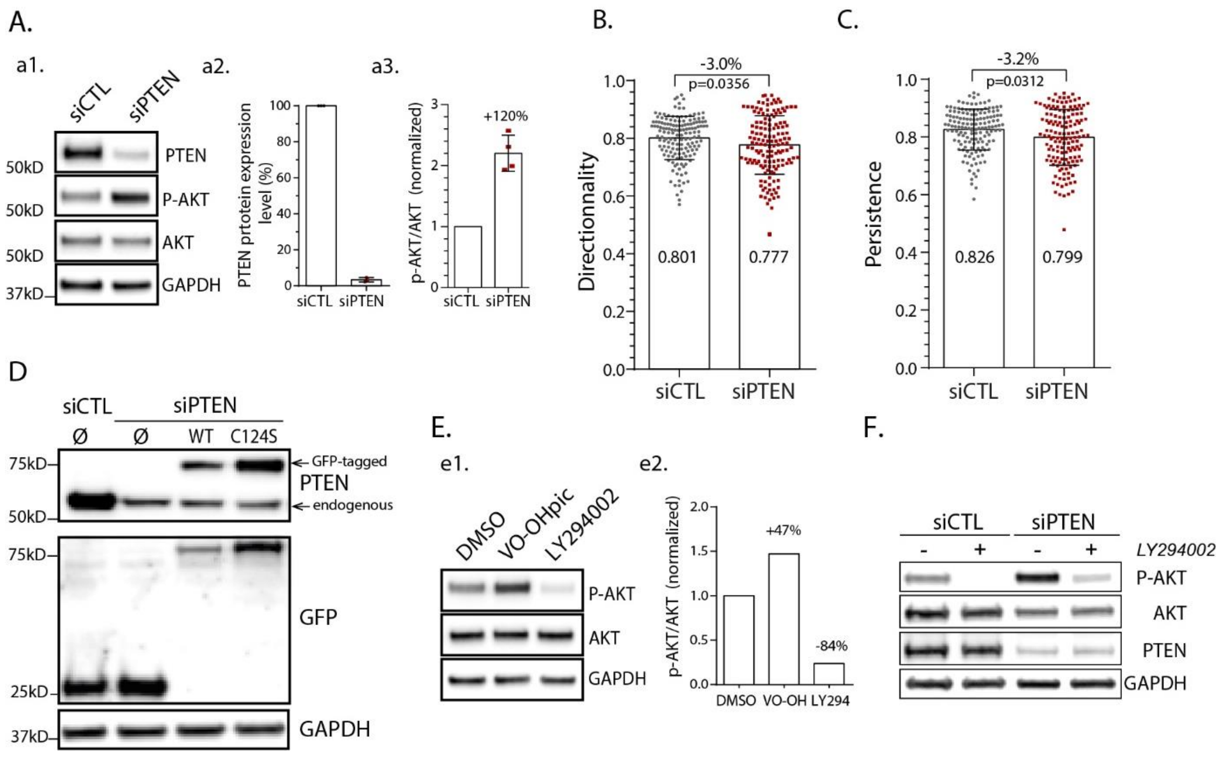


**Supplementary Figure 1. A**. Western Blot analysis of PTEN, phosphor-AKT, AKT and GAPDH in siCTL and siPTEN astrocytes. a2. PTEN protein expression level in siPTEN cells (normalized to siCTL, N=3). a3. Phospho-AKT/AKT ratio level in siPTEN cells increases by 120% compared to siCTL cells. Directionality (**B)** and Persistence (**C**) indexes (see Material and Methods) in siCTL and siPTEN astrocytes migrating in a wound-healing assay (n=150 and 147, N=3, unpaired t-test). **D.** Western Blot analysis of astrocytes infected with lentivirus expressing GFP (Ø), GFP-PTENwt (WT) and GFP-PTEN(C124S) (C124S). Blotting is performed to detect PTEN, GFP and GAPDH. **E.** Western Blot analysis of PTEN, phosphor-AKT, AKT and GAPDH in astrocytes treated with DMSO, VO-OHpic and LY294002. *e2***.** Phospho-AKT/AKT ratio level in drug-treated astrocytes confirms efficiency of the drugs at inhibiting (LY294002) or increasing (VO-OHpic) PI3K signalling. **F.** Western Blot analysis of PTEN, phosphor-AKT, AKT and GAPDH in siCTL and siPTEN astrocytes treated with or without LY294002.

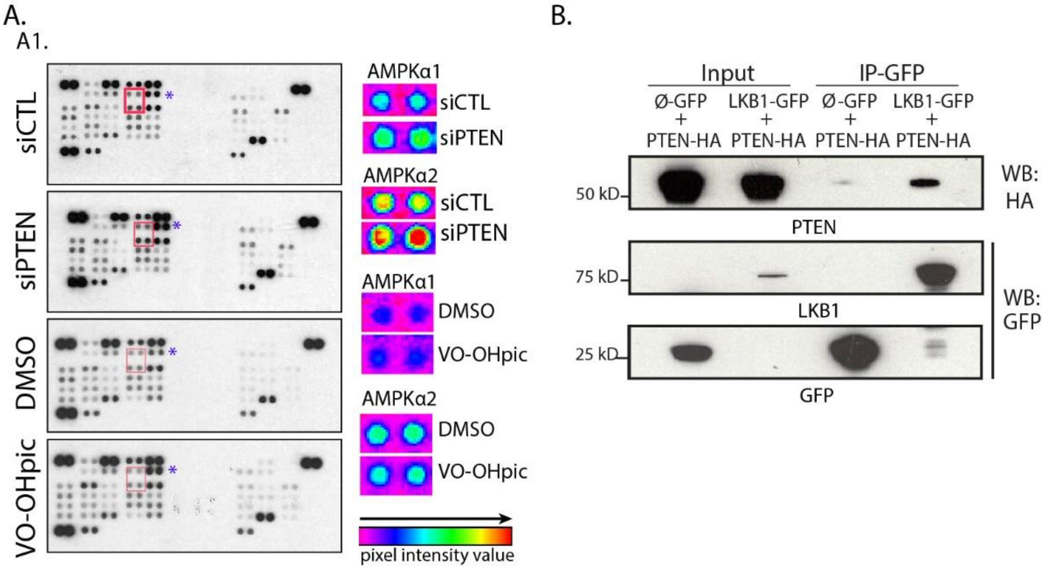


A2.

| | siPTEN/ siCTL | VO-OH/ DMSO |
|---|---|---|
| FAK (Y397) | +10% | -15% |
| **AMPKα1/2 (T172)** | **+17%** | -16% |
| *AKT 1/2/3 (S473) | +37% | +484% |

**Supplementary Figure 2. A.** *a1*. Protein phosphorylation Screen Assay (human Phospho-kinase array kit) reveals PTEN targets affected by PTEN depletion (siPTEN) but not by PTEN lipid phosphatase inhibition (VO-OH). 41 biotinylated phospho-antibodies spotted in duplicate on nitrocellulose membranes are visualized using chemiluminescent detection reagents. Red boxes highlight phospho-AMPKα1/2 signals, which are zoomed in and transformed into multicolour range pixel intensities. Blue asterisks highlight pAKT as a positive control of the assay. *a2*. Table representing FAK, AMPKα and AKT phosphorylation ratios siPTEN/siCTL and VO-OH/DMSO. AMPKα phosphorylation is only seen when PTEN is depleted, not when its lipid phosphatase activity is inhibited. This is the opposite to AKT phosphorylation and similar to FAK phosphorylation, a known PTEN protein phosphatase target. **B**. Co-immunoprecipitation assays in HEK-293T cells transfected with LKB1-GFP and PTEN-HA shows interaction between LKB1 and PTEN.

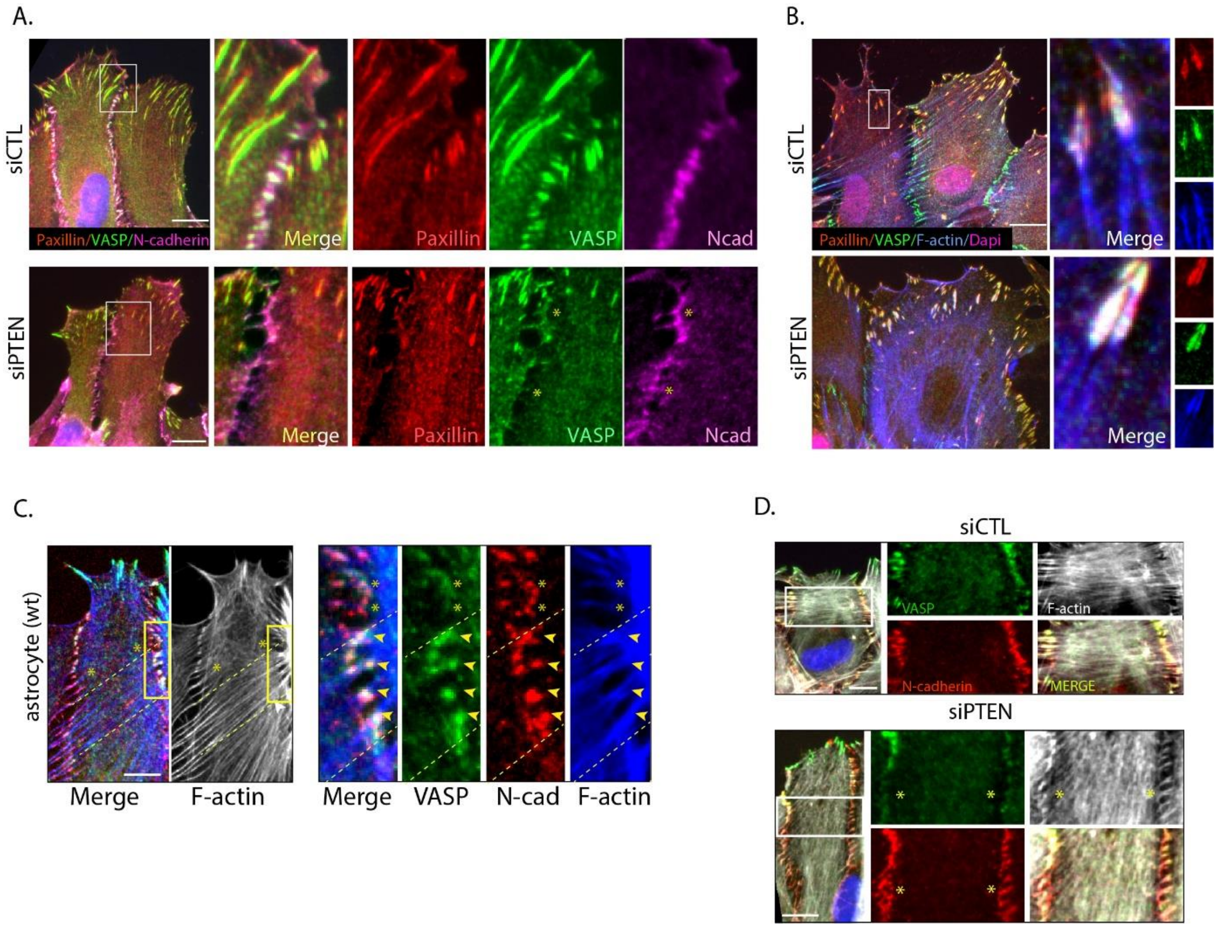


**Supplementary Figure 3. A**. Immunofluorescence images of VASP (green), Paxillin (red), N-cadherin (magenta) and nucleus (blue) in migrating siCTL and siPTEN astrocytes. Boxed region highlight recruitment of VASP both at cell-cell junctions (white pixels in the merge channel) and at focal adhesions (yellow pixels in the merge channel) during cell migration. Note that in siPTEN cells, VASP localises with Paxillin at focal adhesions but not with N-cadherin at cell-cell junctions (yellow asterisks). **B.** Immunofluorescence images of VASP (green), Paxillin (red), F-actin (blue) and nucleus (magenta) in migrating siCTL and siPTEN astrocytes. Boxed region highlight two focal adhesions at the cell front where VASP colocalizes with paxillin and F-actin. No change is observed in siPTEN cells. **C**. Immunofluorescence images of VASP (green), N-cadherin (red), F-actin (blue or gray) in migrating control astrocytes. Yellow dashed lines delineate a zone where interjunctional transverse actin arcs (ITA) are visible and where VASP signal is high at cell-cell junctions. Yellow asterisks highlight a zone where no ITA is visible and where VASP signal is absent from cell-cell junctions. **D.** Immunofluorescence images of VASP (green), N-cadherin (red) and F-actin (gray) in siCTL and siPTEN migrating astrocytes. Note the absence of VASP at cell-cell junctions (yellow asterisks) associated with absence of ITA in siPTEN cells. Scale bar: 10µm.

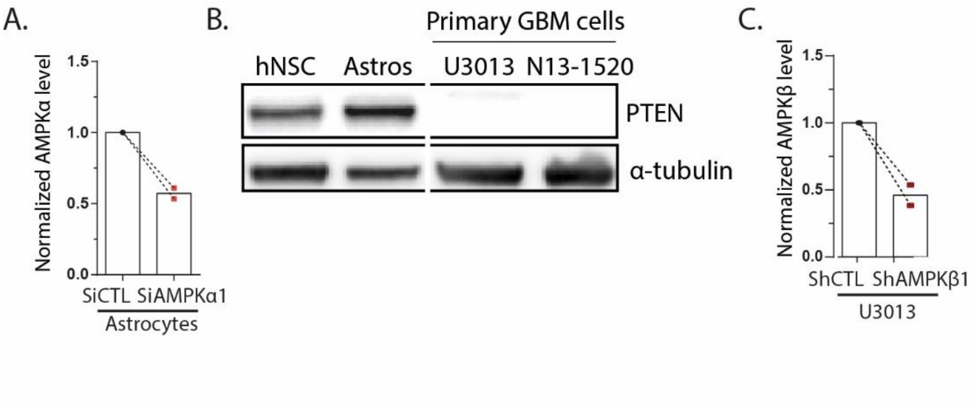


**Supplementary Figure 4. A**. Normalized AMPKα level in siCTL and siAMPKα1 astrocytes from western Blot siRNA efficiency experiments (N=2). **B.** Western Blot showing PTEN and α-tubulin levels in human Neural Stem Cells (hNSC), rat astrocytes (Astros), U3013 and N13-1520 primary GBM cells. **C.** Normalized AMPKβ1 level in shCTL and shAMPKβ1 U3013 GBM cells, from western Blot shRNA efficiency experiments (N=2).

**Movie1: PTEN depletion increases collective glial cell migration velocity**

siPTEN astrocytes close artificial 2D wound faster than siCTL cells. Total time: 22hours, 1 image every 15min. Scale bar: 100µM.

**Movie2: AMPK depletion decreases collective cell migration velocity of siPTEN cells**

siCTL, siPTEN, siAMPKα1 and siPTEN+siAMKPKα1 glial cells closing artificial 2D wounds. Total time: 21hours, 1 image every hour. Scale bar: 100µM.

**Movie3: AMPK inhibition halts PTENneg primary glioblastoma cell invasion**

U3013 cells are embedded in Matrigel® as spheroids and treated with DMSO (left) and Compound C (CC, right). Note that CC treatment prevents the radial invasion of Matrigel observed in DMSO-treated GBM cells. Total time: 24.5hours, 1 image every 15min. Scale bar: 100µM.
